# Metagenomic analysis with strain-level resolution reveals fine-scale variation in the human pregnancy microbiome

**DOI:** 10.1101/266700

**Authors:** Daniela S. Aliaga Goltsman, Christine L. Sun, Diana M. Proctor, Daniel B. DiGiulio, Anna Robaczewska, Brian C. Thomas, Gary M. Shaw, David K. Stevenson, Susan P. Holmes, Jillian F. Banfield, David A. Relman

## Abstract

Recent studies suggest that the microbiome has an impact on gestational health and outcome. However, characterization of the pregnancy-associated microbiome has largely relied on 16S rRNA gene amplicon-based surveys. Here, we describe an assembly-driven, metagenomics-based, longitudinal study of the vaginal, gut, and oral microbiomes in 292 samples from ten subjects sampled every three weeks throughout pregnancy. 1.53 Gb of non-human sequence was assembled into scaffolds, and functional genes were predicted for gene-and pathway-based analyses. Vaginal assemblies were binned into 97 draft quality genomes. Redundancy analysis (RDA) of microbial community composition at all three body sites revealed gestational age to be a significant source of variation in patterns of gene abundance. In addition, health complications were associated with variation in community functional gene composition in the mouth and gut. The diversity of *Lactobacillus iners*-dominated communities in the vagina, unlike most other vaginal community types, significantly increased with gestational age. The genomes of co-occurring *Gardnerella vaginalis* strains with predicted distinct functions were recovered in samples from two subjects. In seven subjects, gut samples contained strains of the same *Lactobacillus* species that dominated the vaginal community of that same subject, and not other *Lactobacillus* species; however, these within-host strains were divergent. CRISPR spacer analysis suggested shared phage and plasmid populations across body sites and individuals. This work underscores the dynamic behavior of the microbiome during pregnancy and suggests the potential importance of understanding the sources of this behavior for fetal development and gestational outcome.

## INTRODUCTION

The importance of the microbiota in human nutrition, immune function, and physiology is well known (Fujimura et al. 2010; Garrett et al. 2010; Charbonneau et al. 2016), and disturbance (such as antibiotic use and disease) has important effects on the microbiome (Cho and Blaser; Costello et al. 2012). Pregnancy is a natural disturbance that can be understood as a special immune state. During gestation, significant hormonal, physiological, and immunological changes allow for, and promote the growth of a developing fetus (Nuriel-Ohayon et al. 2016). For example, some of the changes that occur in pregnancy resemble metabolic syndrome (weight gain, glucose intolerance, and low-level inflammation, among other symptoms). Given the roles of the microbiome during other states of health, it is presumed to provide fundamental support for fetal development as well (Charbonneau et al. 2016). Despite this, few studies of the human microbiome in pregnancy, beyond community composition-based analyses, have been reported. One study investigating the gut microbiota at one time point during each of the first and third trimesters suggested that community composition changes during pregnancy (Koren et al. 2012). Similarly, abundances of viable counts of common oral bacteria have been found to be altered during gestation, especially during early pregnancy, compared to non-pregnant women (Nuriel-Ohayon et al. 2016). However, a detailed longitudinal 16S ribosomal RNA (rRNA) gene survey of the gut and oral microbiota from 49 women during pregnancy demonstrated relative stability over time (DiGiulio et al. 2015).

The vaginal bacterial community composition of pregnant and non-pregnant individuals has been classified into community state types (CST) based on the presence and relative abundance of organisms characterized with the 16S rRNA gene (Ravel et al. 2011; Hickey et al. 2012; DiGiulio et al. 2015). More recently, an alternative classification for vaginal communities was proposed based on the frequency of *Lactobacillus iners, Lactobacillus crispatus*, and *Gardnerella vaginalis* (Callahan et al. 2017). Studies of vaginal community composition based on 16S rRNA gene amplicon sequence analysis in pregnant and non-pregnant individuals have reported relative stability throughout gestation, although transitions between CSTs within subjects have been observed (Gajer et al. 2012; Romero et al. 2014; DiGiulio et al. 2015; McIntyre et al. 2015; Stout et al. 2017). In addition, studies have established associations between the vaginal microbiota and race/ethnicity (Hillier et al. 1995; Zhou et al. 2007; Ravel et al. 2011; Human Microbiome Project 2012; Hyman et al. 2014) and, more recently, associations with premature birth (Callahan et al. 2017; Kindinger et al. 2017; Stout et al. 2017; Tabatabaei et al. 2018).

Despite this recent attention to the microbiome during pregnancy, relatively few studies have discriminated among bacterial strains, and few have described the phage populations in the human microbiome during pregnancy. Since CRISPR spacers are mainly derived from phage, plasmids, and other mobile elements (reviewed extensively in Horvath and Barrangou 2010; Karginov and Hannon 2010; Marraffini and Sontheimer 2010), the CRISPR-Cas system can be used to identify these sequences (Bolotin et al. 2005; Mojica et al. 2005; Pourcel et al. 2005; Andersson and Banfield 2008; Sun et al. 2016) where spacers reflect exposures to phage and plasmids in a whole community metagenomic dataset.

The goal of this work was to survey the genome content and functional potential of microbial communities during gestation. We describe the vaginal, gut and oral microbiomes from ten pregnant subjects from first trimester through delivery, with the goal of broadening our understanding of the functional potential and the dynamics of the human microbiome during pregnancy.

## RESULTS

### Community structure from assembly-driven metagenomics

Ten pregnant women whose microbiota have been studied previously using 16S rRNA gene amplicon data (DiGiulio et al. 2015) were selected for further gene and genome composition analysis. Subjects were selected in part based on their compliance in providing samples on a regular basis from multiple body sites. Six subjects delivered at term and four delivered preterm (i.e., <37 weeks gestation). Five subjects (four who delivered at term, and one who delivered preterm) were diagnosed with having some type of pregnancy complication: preeclampsia, Type II diabetes, and oligohydramnios (Supplemental Table S1).

Shotgun metagenomic sequencing was performed on 292 vaginal, oral, and gut samples that had been collected, on average, every three weeks over the course of gestation (Table 1; Supplemental Table S1). Community composition and estimates of diversity were evaluated from full-length 16S rRNA gene sequences that were reconstructed from metagenomics reads using EMIRGE (Dick et al 2009), and from the average number of single copy ribosomal protein (RP) sets encoded in assembled scaffolds. Based on reconstructed 16S rRNA gene sequences, 1,553 taxa (clustered at 97% identity) were identified. In total, 22,737 predicted proteins annotated as one of the 16 RPs described for phylogenetic classification in Hug et al.(Hug et al. 2016) were encoded in 4,650 assembled scaffolds at all body sites. The RP sets were clustered at 99% average amino acid identity (AAI) into 4,024 taxa (see Methods) (Table 1).

**Table 1.** Metagenomic sequencing statistics. Stool and rectal swab sequences were combined during co-assembly per subject. Average richness* (genome types) was estimated from the average number of single-copy ribosomal protein sets. M, million. The average number of reads per sample (M) was calculated prior to removal of human contamination. The average number of reads per assembly (M) was calculated from the average number of reads per sample that mapped to the co-assemblies.

|  | Vagina | Gut |  | Saliva |
| --- | --- | --- | --- | --- |
|  |  | Stool | Rectal |  |
| Samples analyzed | 101 | 34 | 56 | 101 |
| Reads sequenced (M) | 3478 | 609 | 1212 | 1903 |
| Avg. reads per sample (M) | 34 | 18 | 22 | 19 |
| Avg. human contamination | 97% | 32% | 42% | 82% |
| Avg. reads per assembly (M) | 7.89 | 117.6 |  | 27.19 |
| Bases assembled, >1 Kb (M) | 177 | 2900 |  | 1345 |
| GC (%) range | 25 - 63 | 25- 61 |  | 28 - 66 |
| Max coverage | 1175 | 996 |  | 138 |
| Average richness* | 9 ($\pm 2$) | 101 ($\pm 15$) | | 42 ($\pm 11$) |
| ORFs predicted (M) | 0.19 | 3.13 |  | 1.60 |

Estimates of richness and evenness from reconstructed 16S rRNA genes differed depending on body site: vaginal communities were the least diverse and gut communities were the most diverse (Supplemental Fig. S1A). Vaginal communities appeared to be largely under-sampled with the metagenomic data in comparison to the previously-published 16S rDNA amplicon data (Digiulio et al. 2015), perhaps due to high levels of human sequence contamination in the metagenomic data (Supplemental Fig. S1B, vagina). It is also possible that the amplification of 16S rDNA leads to higher richness than expected. By contrast, gut communities showed higher richness and evenness (visualized as the slope of the curves) when viewed with metagenomics than with 16S rRNA gene surveys (Supplemental Fig. S1B, gut) possibly due to higher resolution from deep sequencing in the metagenomics data, and/or from the so-called ‘universal’ 16S rRNA gene primers not amplifying all microbial targets. On the other hand, the richness of the oral communities recovered from metagenomics appeared to approximate that observed in 16S rDNA amplicon (Supplemental Fig. S1B, saliva).

Unlike gut or salivary communities, vaginal communities were generally dominated by one organism based on whole genome abundance measures (Fig. 1A) and reconstructed 16S rRNA gene analysis (Supplemental Fig. S2A). The taxonomic classifications, community state types (CSTs), and community dynamics revealed through analysis of 16S rRNA gene sequences assembled by EMIRGE as well as from ribosomal protein (RP) sequences, resembled those revealed by 16S rRNA gene amplicon sequence data (DiGiulio et al. 2015) (Supplemental Fig. S2). Communities dominated by *L. iners* showed a clear increase in taxonomic richness towards the end of pregnancy (subjects Term2, Term3, Term4, and Pre4), while those dominated by *L. crispatus* appeared to remain stable over time (subjects Term5, Term6, Pre1, and Pre3) (Fig. 1A). To explore the observed association between increasing community richness and community domination by *L. iners*, we re-analyzed the 16S rRNA amplicon data reported previously (DiGiulio et al. 2015; Callahan et al. 2017) (Supplemental Fig. S3). Shannon’s diversity significantly increased with gestational age in *L. iners*-and *L. crispatus*-dominated communities in the Stanford population (F = 32.07, p < 0.0001; and F = 8.32, p = 0.004; respectively) but not in the UAB population. When *L. iners*-and *L. crispatus*-dominated communities were each pooled from both populations, gestational age trends achieved significance in both cases (*L. iners*, F = 23.23, p < 0.0001; *L. crispatus*, F = 8.61, p < 0.0034). These data suggest dynamic vaginal community composition over the course of gestation.

**Figure 1.**
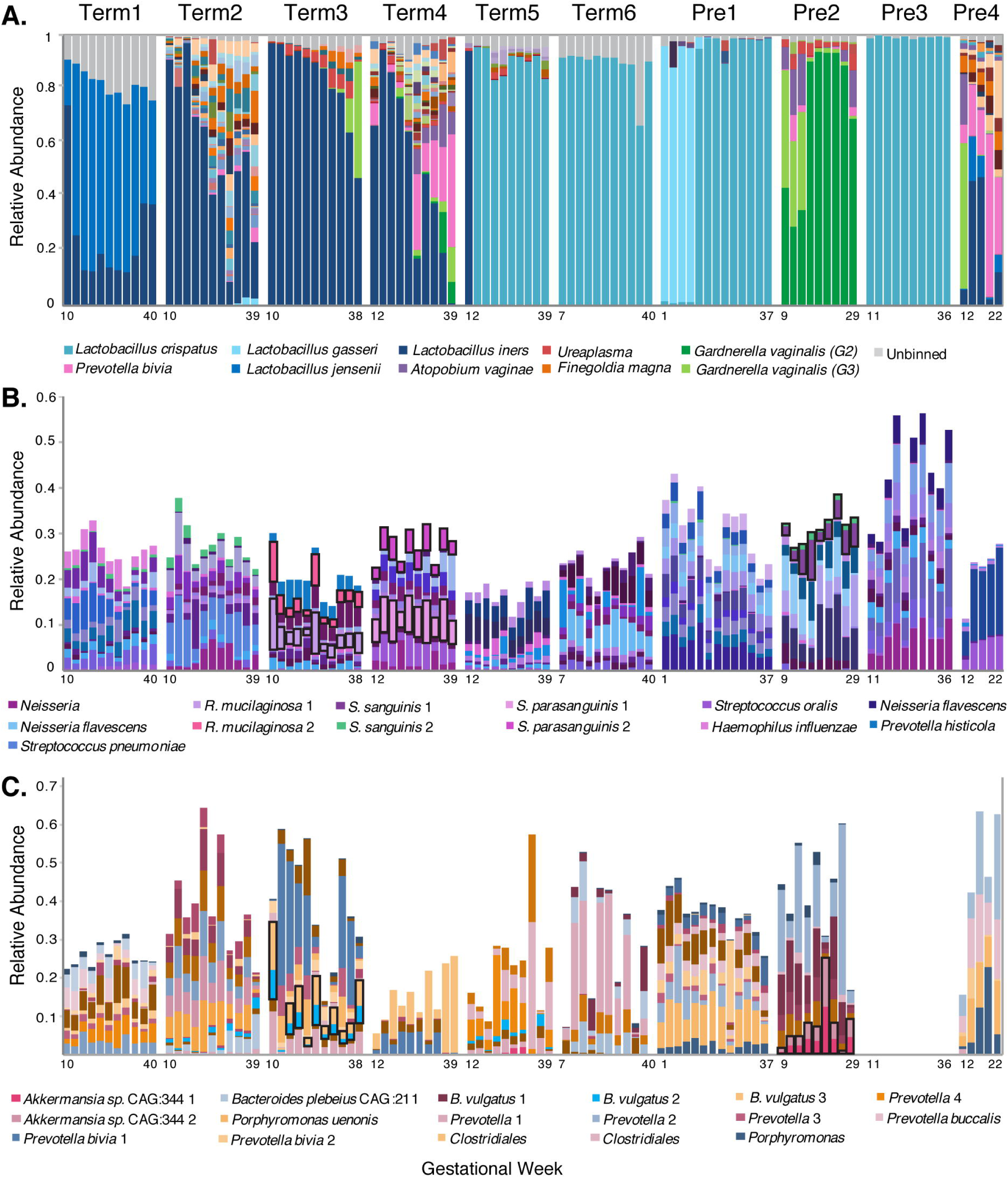
Community structure of the pregnancy microbiome over time in 10 subjects. **A.** Relative abundance of vaginal genome bins (y axis). Abundance was estimated from the number of reads that mapped to each bin and normalized by the length of the bin. The top 10 most abundant vaginal taxa are displayed (see key at bottom). Unbinned: sequences that were not assigned to a classified genome bin. **B. and C.** Relative abundance (y axis) of the top 50 most abundant taxa across all subjects in saliva (B) and gut (C) samples, respectively. Gut samples from subject Pre3 were not available. Species abundance was estimated from the average read counts of single-copy ribosomal protein (RP) sets (at least one of 16), summed over scaffolds sharing RPs clustered at 99% amino acid identity. Each taxon is represented by a distinct color (see key for selected taxa at bottom) and is classified at the most resolved level possible. Co-occurring strains of *Rothia mucilaginosa, Streptococcus sanguinis,* and *Streptococcus parasanguinis* in (B), and of *Bacteroides vulgatus,* and *Akkermansia* spp. CAG:344 in (C) are highlighted with black boxes.

Due to the high diversity of gut and saliva communities, the relative abundances of 16 single-copy ribosomal protein (RP) sets were used to estimate the dynamics of gut and saliva communities over time. The relative abundances of the most abundant taxa in the gut and saliva showed relative stability over time within subjects but high inter-individual variability (Figs. 1B and 1C). For example, the 9 most abundant taxa in the saliva samples of subject Term5 were relatively consistent in abundance over all time points, representing at most 20% of the community within that subject, but were distinct from the most abundant taxa in the saliva of the other subjects (Fig. 1B). In addition, strains could be resolved from RP cluster variation within scaffolds, and co-occurring bacterial strains were observed in gut and saliva samples within subjects (Fig. 1B and 1C, black boxes).

Most taxa identified by 16S rRNA gene reconstruction (classified at 97% average nucleotide identity, or ANI) and RP sets (classified at 99% average amino acid identity, or AAI) were shared between body sites among all subjects, but the level of sharing was most evident between vaginal and gut samples (Supplemental Fig. S1C). Phyla found uniquely in gut samples included the *Bacteria*, Verrucomicrobia and Synergistetes, the *Archaea,* Euryarchaeaota, and the *Eukarya,* Stramenopiles (*Blastocystis homini*); whereas phyla found uniquely in oral samples included the *Bacteria,* Spirochaetes and SR1. Novel taxa were detected in oral and gut samples: three 16S rRNA sequences in saliva could not be classified to a known phylum and three taxa represented by novel RP sets could not be classified at the domain level based on searches in the public databases. Similarly, six 16S rRNA gene sequences in gut samples could not be classified at the phylum level, and 10 RP sets remained unclassified at the domain level.

### Sources of variation in community-wide gene profiles

To gain insights into all potential sources of variation in the patterns of gene abundance across all subjects and samples, non-metric multidimensional scaling (NMDS) was performed on variance-stabilized abundances of gene families represented in the UniRef90 database. NMDS revealed three major sources of variation in the patterns of gene abundance: subject, gestational age, and health complication. Samples clustered primarily based on subject for each body site (Fig. 2A; Supplemental Figs. S4B and S4C). In addition, when considering vaginal samples, gestational age was a major source of variation for subjects with high diversity communities (Supplemental Fig. S4A, Pre4, Term3, and Term4). Taxonomic assignment was also associated with gene abundance profiles in the vagina. Specifically, gene profiles from *Gardnerella vaginalis* appeared correlated with positive scores on NMDS2 (samples from subject Term4); *L. gasseri, Sneathia* and *Prevotella* gene abundances were associated with positive NMDS1 scores (samples from subject Term2); and *Lactobacillus crispatus, L. iners, and L. jensenii* gene abundances were associated with negative scores along NMDS2 (samples from all other subjects) (Fig. 2B). NMDS plots of variance-stabilized UniRef90 gene family abundances in the distal gut microbiota showed a subtle effect of Shannon’s diversity index on sample variation (Supplemental Fig. S5) and plotting Shannon’s diversity index from ribosomal protein sets, from reconstructed 16S rRNA gene sequences via EMIRGE, and from UniRef90 gene families, indicated a decrease in diversity over time in our 10 subjects (Supplemental Fig. S5). Finally, in oral and gut communities, health complication explained some variation in community gene composition: samples from the 3 subjects who were diagnosed with preeclampsia (two who delivered at term and one preterm) appeared to cluster separately from samples collected from subjects with other health states and pregnancy outcomes (Figs. 2D and 2F).

**Figure 2.**
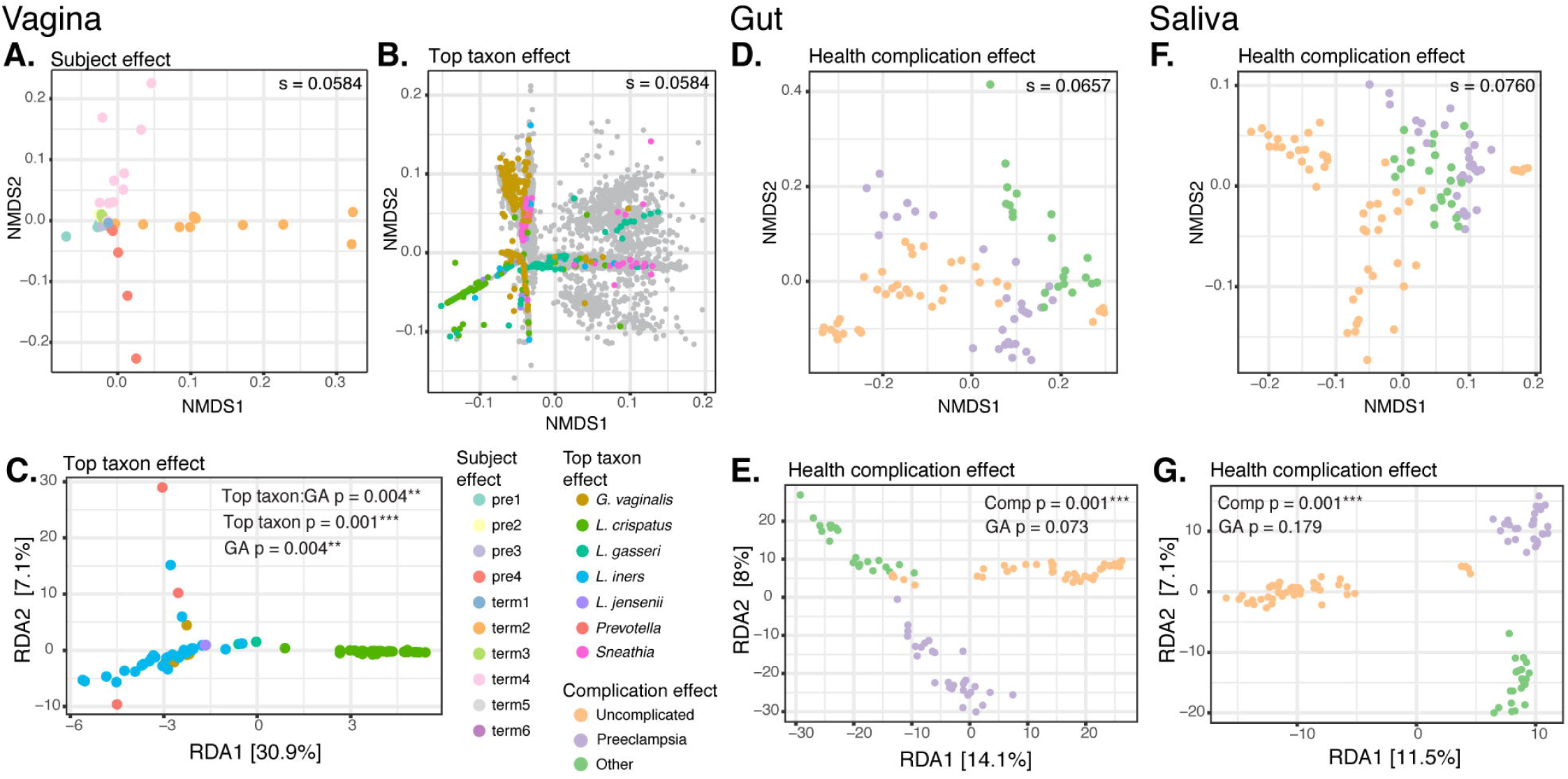
Sources of variation in abundance of UniRef90 gene families across all subjects and samples. Top **(A-D)**: Non-metric multidimensional scaling (NMDS) plots from Bray-Curtis distance matrices of variance-stabilized gene family abundances. The stress, “s” (the amount of variability unexplained by the NMDS ordination) is shown on each plot. Bottom **(E-G)**: Redundancy analysis (RDA) plots from variance-stabilized gene family abundances. The p-values for the RDA plots were estimated with the anova.cca function from the vegan package in R. **A., B.** NMDS split plot from vaginal samples: the “samples” plot (**A**) was color-coded based on subject, while the “genes” plot (**B**) was color-coded based on the taxonomic classification of genes (grey dots: genes belonging to other taxa). **E.** RDA plot from vaginal samples, constrained by gestational age (GA) in weeks and by the most abundant taxon in each sample. Samples are color-coded based on the most abundant taxon. **C., F.** NMDS and RDA plots from gut samples. **D., G.** NMDS and RDA plots from saliva samples. Gestational age and health complication were used to constrain the RDA analysis in **(F)** and **(G)**. Complication: uncomplicated (5 subjects); preeclampsia (3 subjects); other, i.e., type II diabetes (1 subject); and oligohydramnios (1 subject).

To quantify the relative influence of gestational age, dominant taxon (vaginal microbiota), and health complication on community composition, redundancy analysis (RDA) was performed on variance-stabilized gene family abundances. Although the effect of gestational age was significant only for vaginal communities (F = 2.71, p = 0.017) when data from all subjects and samples were considered (Figs. 2C, 2E, and 2G), gestational age trends were significant at all body sites and for most subjects, and showed stronger effects, when the data for each subject were examined separately (Fig. 3; Supplemental Fig. S6). For vaginal samples, significant trends with gestational age were observed in 6 of 10 subjects (subjects Pre1, Pre2, Pre3, Term2, Term3, Term4) even in low-diversity communities dominated by *L. crispatus* (subjects Pre1 and Pre3) (Supplemental Fig. S6A). The most abundant taxon in vaginal communities, and health complication (preeclampsia, diabetes, oligohydramnios, or uncomplicated pregnancy) in gut and oral communities were highly significant sources of variation in gene family abundances based on RDA of the overall data (vagina: F = 14.97, p = 0.001; saliva: F = 11.20, p = 0.001; gut: F = 12.34, p = 0.001) (Figs. 2C, 2E, and 2G), although the numbers of subjects with each of these complications were small.

**Figure 3.**
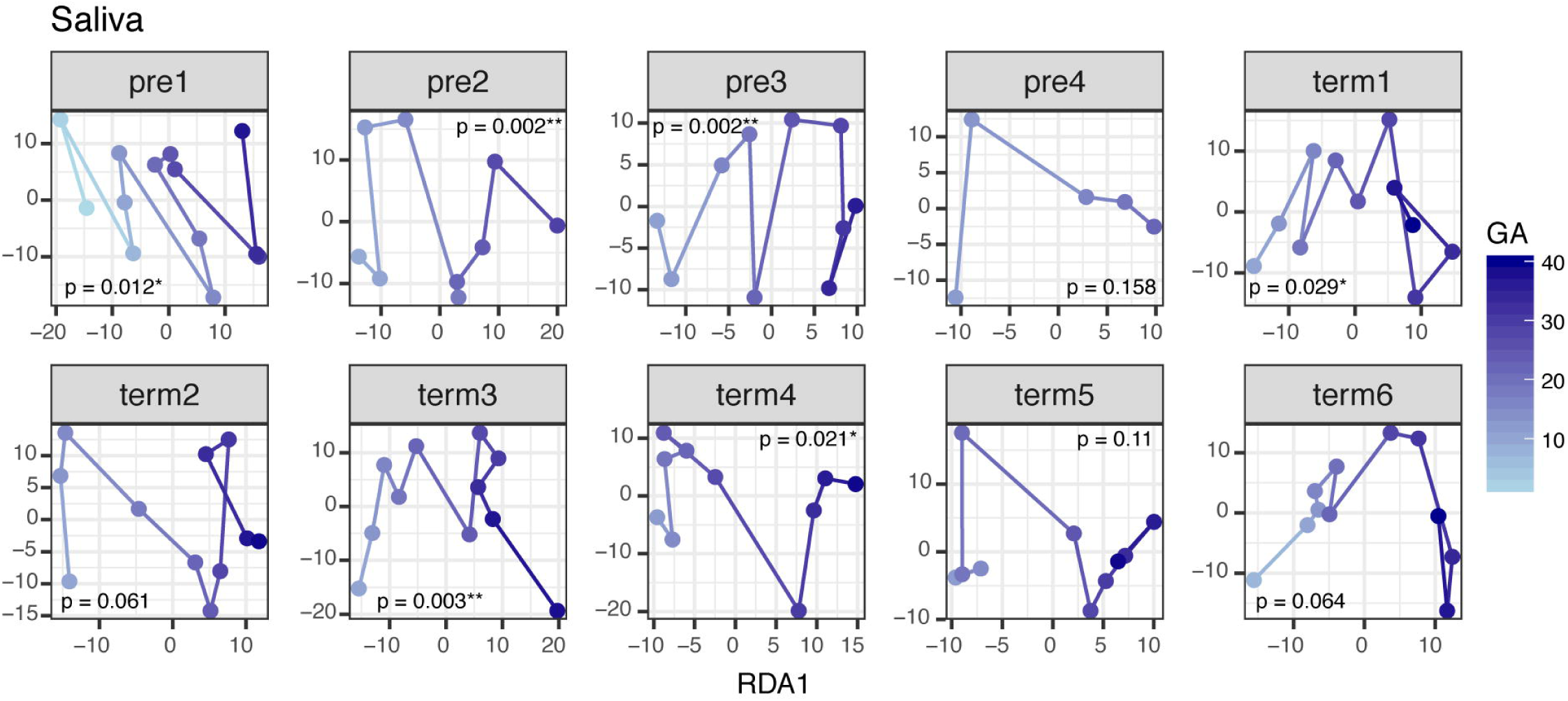
Gestational age trends for abundances of gene families in saliva samples for each subject. Gestational age (GA) was used to constrain the redundancy analysis (RDA) of variance-stabilized gene family abundances within individuals. Gestational age effect is observed along the x-axis (RDA1 axis), and points within plots were connected based on the resulting ordination scores. P-values were calculated with anova on the RDA ordination constraint using the anova.cca function of the vegan package in R.

Overall pathway composition and abundance remained stable over time at all body sites when all subjects were viewed together (Supplemental Fig. S7, top plots). Not surprisingly, core metabolic pathways consistently accounted for most of the overall pathway composition and abundance in all subjects. When pathways were viewed individually, their abundances were stable over time (e.g., glycolysis at all body sites, Supplemental Fig. S7, bottom plots); however, some pathways were variable over gestational age. For example, the relative abundance of the enterobactin biosynthesis pathway (a siderophore-mediated iron uptake system (Raymond et al. 2003)) decreased towards the end of pregnancy in most gut and oral samples. Yet, the enterobactin biosynthesis pathway identified in some samples remain stable at low basal levels (ENTBACSYN, Supplemental Fig. S7, Gut, bottom plots). These results suggest that iron may have been in plentiful supply as pregnancy progressed in the oral and gut sites, leading to enrichment of organisms without enterobactin biosynthesis capabilities. In addition, the increasing relative abundance of the pyruvate fermentation to acetate and lactate pathway towards the end of pregnancy in gut samples (PWY-5100, Supplemental Fig. S7, Gut) suggests that fermenters might become enriched in the gut during pregnancy.

### Gardnerella vaginalis strains co-occur in the vagina

Gardnerella vaginalis is an important species associated with both bacterial vaginosis (Schwebke et al. 2014) and an increased risk of preterm birth (Callahan et al. 2017). The near-complete draft genomes of six strains of *G. vaginalis* from the vaginal samples of four subjects were recovered in this study (Fig. 1A, colored light and dark green). *G. vaginalis* genomes were abundant early in pregnancy in two subjects who delivered preterm (Pre2 and Pre4), and they dominated samples later in pregnancy in two subjects who delivered at term (Term3 and Term4). Two subjects carried co-occurring *G. vaginalis* strains (Figs. 1A and 4A; subjects Term4 and Pre2). The strains shared >99% average pairwise nucleotide identity in the 16S rRNA gene yet single nucleotide polymorphism (SNP) patterns were evident in this gene (Supplemental Fig. S8A). One strain was generally more abundant than the other throughout pregnancy: in subject Term4, both *G. vaginalis* strains were found at low relative abundance (less than 7% for each strain in any sample) (Fig. 4A); while in subject Pre2, the two strains co-existed at roughly equal abundances early in pregnancy and then one assumed dominance as gestation progressed (Fig. 4A).

**Figure 4.**
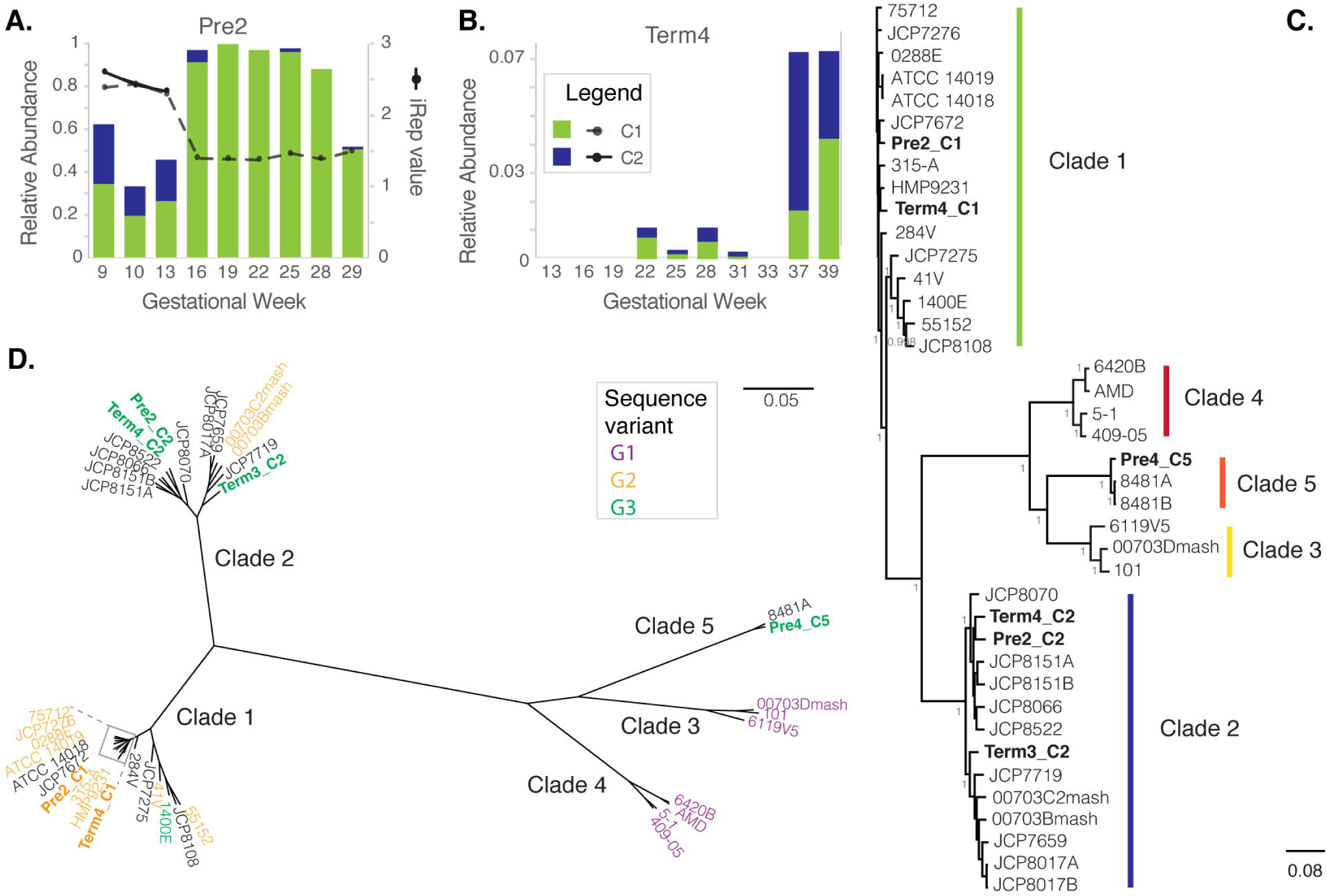
*Gardnerella vaginalis* genome analysis. 16S rRNA gene abundance of *G. vaginalis* strains recovered from each of two subjects, Pre2 **(A)** and Term4 **(B)**. *G. vaginalis* genotypic groups, C1 and C2 are colored according to the classification in **(C)**. Relative abundance (left y-axis). Estimated iRep values are plotted for *G. vaginalis* strains in subject Pre2 (right y-axis). **C.** Phylogenomic tree of 40 *G. vaginalis* strains genomes, including 34 available in GenBank. The 6 genomes recovered in the current study are shown in bold. Colored bars represent genotypic groups within the *G. vaginalis* phylogeny, where colors for clades 1-4 match the *G. vaginalis* genotypic groups defined by Ahmed *et al* (2012). `Fastree` branch support for the most visible nodes is shown. **D.** Radial representation of the same phylogenomic tree displayed in **(C)**, where leaves are colored based on 16S rRNA V4 sequence variant classification defined by Callahan *et al*. (2017). Genomes for which a full-length 16S rRNA sequence or V4 sequence were not available are shown in black.

To determine whether replication may have been associated with the relative abundance differences in *G. vaginalis* strains within each subject, iRep values were determined. iRep values have been employed to infer replication rates of microbial genome types from draft quality genomes reconstructed from metagenomic data and can be used to suggest that a particular microbial type is actively growing at the time of sampling (Brown et al. 2015). As described in (Brown et al. 2015), an iRep value of 2 would indicate that most of the population is replicating 1 copy of its chromosome (Brown et al. 2015). The iRep values for the co-occurring *G. vaginalis* strains in subject Pre2 indicate both strains were actively replicating at the beginning of pregnancy (iRep ∼2.5) when both strains co-existed at relatively equal abundances. After gestational week 16, the Pre2_C1 strain remained dominant with a stable replication rate value of ∼1.5 (that is, roughly 50% of the population replicating) while the Pre2_C2 strain dropped below iRep calculation limits due to low abundance (Fig. 4A).

A full-length multiple genome alignment and phylogenomic tree of the six genomes recovered in this study, and 34 complete and draft genomes available in GenBank revealed five divergent clades (Fig. 4B,C). The dominant genomes recovered from subjects Term4 and Pre2 (Term4_C1, and Pre2_C1) were more similar to those of a similar genome type found in other subjects than to the co-occurring genome within the same subject (Fig. 4B). When using the ATCC 14019 type strain genome as reference, the total number of SNPs across the genomes did not correlate with the group phylogeny. In other words, a divergent clade does not necessarily harbor more SNPs than strains within Clade 1 to which the ATCC 14019 strain belongs (SNP counts are displayed next to the genome ID in Supplemental Fig. S8B).

From over 50,600 genes predicted in all 40 strain genomes, ∼4,000 genes were unique to a genome, and the rest could be clustered at >60% AAI over 70% alignment coverage into >3,300 orthologous groups. In total, 41,245 orthologous genes were present in at least half the genomes within each of the five clades described above (1763 orthologous groups) and ∼44% of these represented the pangenome (cluster 4, Supplemental Fig. S9A). Furthermore, hierarchical clustering suggested groups of orthologous genes specific to each genome clade: genes shared within Clade 1 genomes (cluster 6), genes in Clade 2 genomes (cluster 13), genes in Clade 3 genomes (cluster 1), genes in Clade 4 genomes (cluster 3), genes in Clade 5 genomes (cluster 5), and one cluster composed of genes shared between Clade 3, Clade 4, and Clade 5 genomes (Supplemental Fig. S9A). A survey of the functional categories represented within these clusters indicated that although there is redundancy of functions across all clusters (most of them also represented in the pangenome cluster) some functions are enriched in clade-specific genomes. For example, hypothetical proteins are enriched in Clade 5 genomes; proteins involved in vitamin and cofactor metabolism, and CRISPR Cas genes are enriched in Clade 3 genomes; and membrane proteins, DNA recombination/repair proteins, and proteins with transferase activity appear enriched in Clade 2 genomes. The results suggest a clade-specific enrichment of functional categories in *G. vaginalis* genomes.

A recent detailed survey of SNP patterns in the V4 region of the 16S rRNA gene from a longitudinal study of vaginal communities from pregnant women classified *G. vaginalis* strains into amplicon sequence variants (ASVs) (Callahan et al. 2017), and the three most frequent sequence variants could be mapped onto divergent clades on a phylogenomic tree (Fig. 4D). The genomes and full-length 16S rRNA gene of two of the described ASVs were recovered in this study: variant G2 (Term4_C1 and Pre2_C1) (tree leaves colored in orange, Fig. 4C); and variant G3 (Term3_C2, Term4_C2, and Pre4_C5), which intermingle across the phylogenomic tree (tree leaves colored in green, Fig. 4C).

### Lactobacillus genomes vary in their mobile elements

Vaginal bacterial communities of many women are dominated by *Lactobacillus iners* (Ravel et al. 2011; DiGiulio et al. 2015), an organism that has been associated with both health and disease (Macklaim et al. 2011). Recent studies have shown that *L. iners* is associated with high diversity CSTs while *L. crispatus* (dominant species in CST I) is thought to be more protective against disease (Callahan et al. 2017; Smith and Ravel 2017). We recovered 6 near complete *L. iners* genomes, as well as 4 fragmented *L. crispatus* genomes (completeness was assessed based on single copy genes and average genome size).

A comparison of the *L. iners* genomes with the KEGG Automatic Annotation Server (Moriya et al. 2007) revealed no major differences in terms of the presence or absence of genes of known function. Alignment of the consensus *L. iners* genome sequences from five of the six strains to the NCBI reference strain DSM 13335 using Mauve showed that the genomes share high levels of synteny (the genome from Pre4 was not used due to its high fragmentation) (see Methods; Fig. 5A). Primary differences between strains were the variable presence of mobile element-like genes including classic phage and plasmid genes (such as phage integrases, phage lysins, phage portal genes, as well as genes putatively involved in restriction-modification systems, toxin-antitoxin systems, and antibiotic transport), and many hypothetical genes (Fig. 5A, islands). Analysis of orthologous gene groups among the six genomes assembled here and 18 other publicly available genomes confirmed that *L. iners* genomes are highly conserved, sharing 90.5% of their genes (Supplemental Fig. S10).

**Figure 5.**
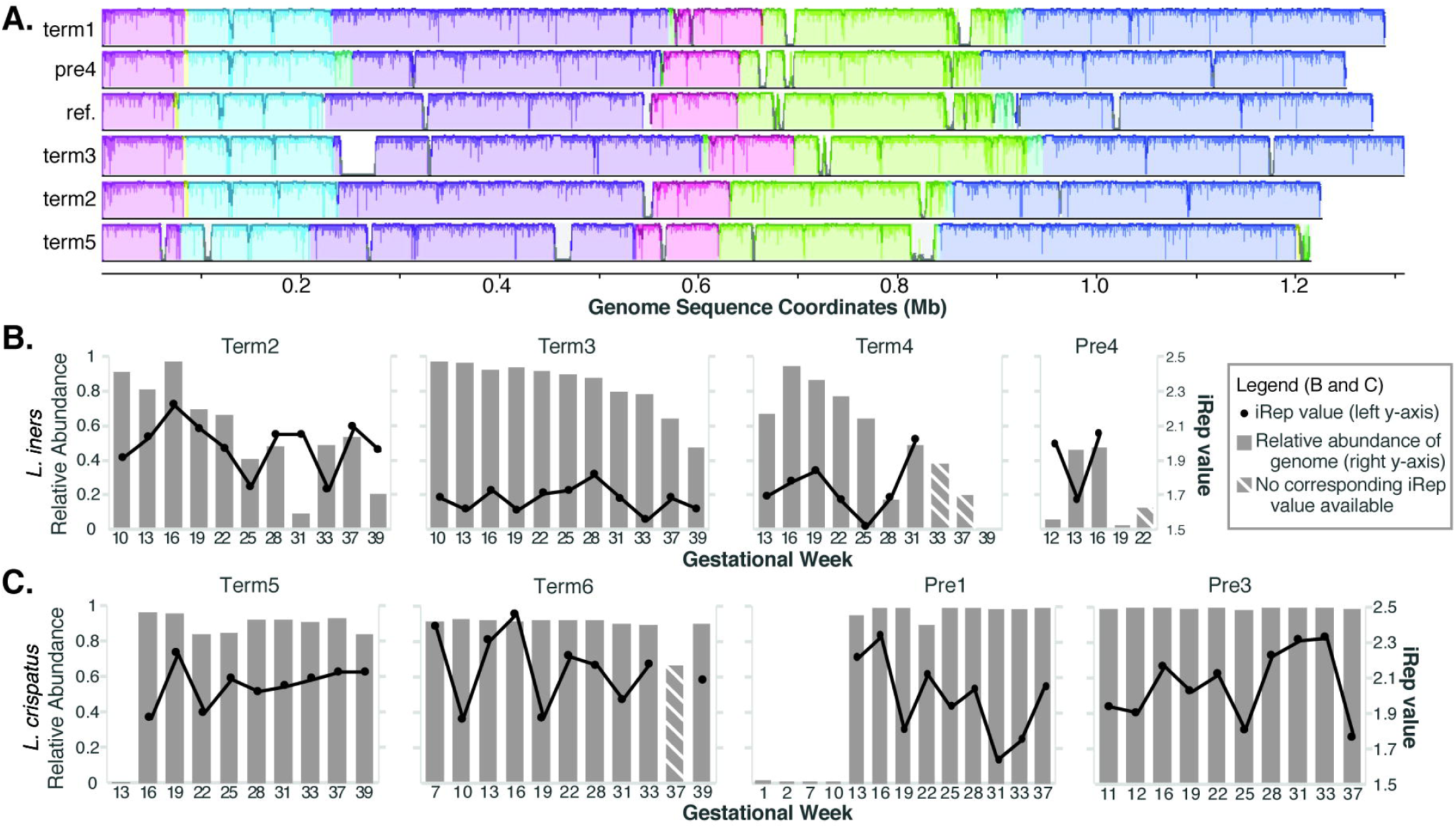
Comparative genomics of *Lactobacillus* spp. *and* estimated replication rates. **A.** Multiple genome alignment of 5 *L. iners* genomes recovered through metagenomics along with the reference strain DSM 13335 (see Methods). A modified alignment created in Mauve shows the shared genomic context, as well as genomic islands unique to a strain (white areas within a conserved block). **B. and C.** Relative abundance and estimated genome replication values (iRep) for *L. iners* and *L. crispatus*, respectively.

The community composition of *L. crispatus*-dominated communities remained generally stable through pregnancy and tended to have low richness (Fig. 1 and Fig. 5C). A full genome alignment and analysis of orthologous genes of 41 draft genomes of *L. crispatus*, including six reconstructed in this study, grouped 80,484 genes into 4,705 orthologous groups present in at least 2 genomes. Hierarchical clustering of the presence/absence of orthologous groups suggested two divergent clusters with more than half of the genes representing the core genome (Supplemental Fig. S11). The two *L. crispatus* types differed in the presence of core SNPs (SNPs present in all genomes) as well as in the presence of cluster-specific orthologous groups. A previous comparative genomics analysis of 10 *L. crispatus* genomes identified ∼1,200 core genes as well as genome-specific genes encoding up to 40% genes of unknown function (Ojala et al. 2014). Characterization of the *L. crispatus* group-specific genes might improve our understanding of the role of this species in human urogenital health.

iRep values were estimated for *L. iners* in subjects Term2, Term3, Term4, and Pre4, and for *L. crispatus* in subjects Term5, Term6, Pre1, and Pre3 to determine whether the different *Lactobacillus* genomes were replicating (see Methods). iRep values fluctuated around 2 (most members of the population were replicating 1 copy of their chromosome) for both organisms and appeared independent of relative abundance (Figs. 5B and 5C). On average, *L. crispatus* had a slightly higher iRep value (2.0) than did *L. iners* (1.8) (∼100% of the *L. crispatus* population, and 80% of the *L. iners* population replicating 1 copy of their genome) (Figs. 5B and 5C).

### CRISPR-Cas diversity and CRISPR spacer targets

In order to investigate the phage and plasmid populations at the different body sites over the course of pregnancy, we characterized the types and distribution of the CRISPR-Cas systems in the vaginal, gut, and saliva assemblies (Fig. 6A). Prior studies have defined several CRISPR-Cas type systems based on the composition and genomic architecture of Cas genes (Makarova et al. 2015; Burstein et al. 2017). Here we identified CRISPR-Cas systems using the five-type classification scheme derived from (Makarova et al. 2015), four of which were identified in our metagenomics data. Type I (thought to be the most common CRISPR-Cas system type (Makarova et al. 2015)) was the dominant CRISPR-Cas system in gut samples, whereas Type II was the dominant system in saliva samples (Fig. 6A).

**Figure 6.**
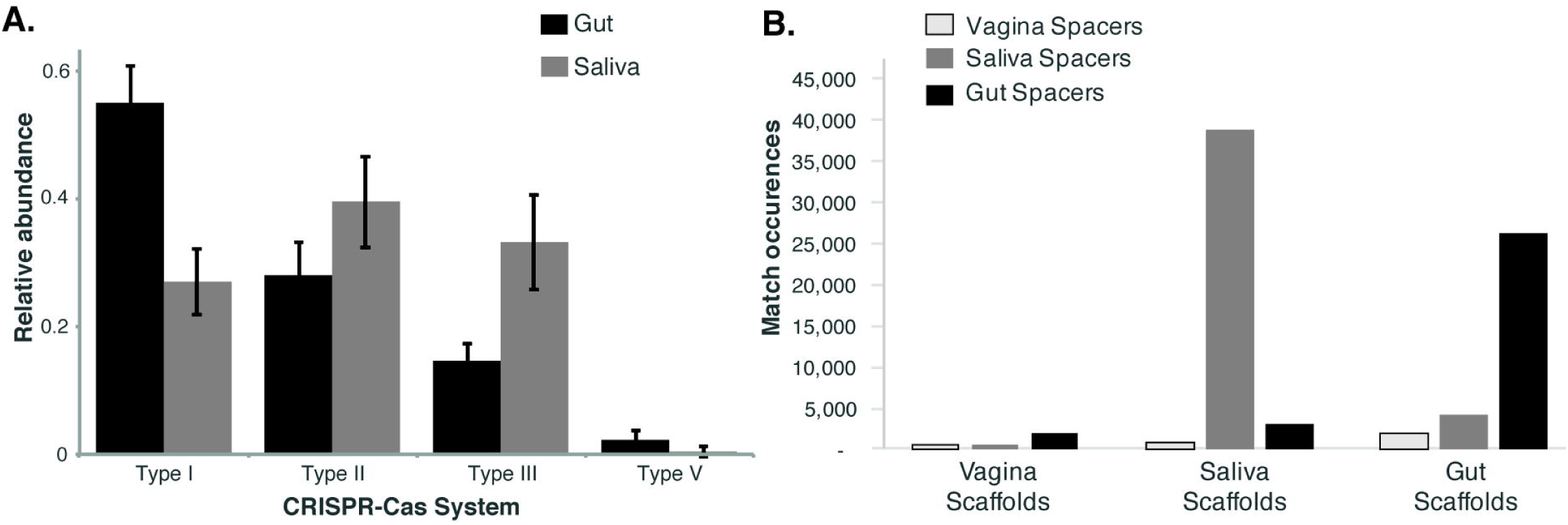
Diversity and distribution of CRISPR-Cas systems and CRISPR spacer targets (mobile elements). **A.** Distribution of CRISPR-Cas system types in the gut (gray) and saliva (black) samples in this study (I, II, III, V). Vaginal samples are not shown due to low bacterial diversity. The black lines indicate standard deviation. **B.** Frequency of matches between spacers and scaffolds. This graph shows the total number of matches by body site (outlined gray, vaginal spacers; light gray, saliva spacers; and dark gray, gut spacers). There were a total of 78,054 matches between spacers and all pregnancy scaffolds. Overall, we detected 3,254 spacer types from vaginal samples, 36,477 spacer types from gut samples, and 36,279 spacer types from saliva samples.

In total, there were 137, 618, and 1255 ‘repeat types’ among vaginal, saliva, and gut samples, respectively (Fig. 6B, Supplemental Table S2). Each repeat type represents a repeat sequence found in a CRISPR locus; different CRISPR loci (within or between organisms) can share the same repeat type. Many more spacer types were found in the gut (36,477) and saliva (36,279) samples than in the vaginal samples (3,254). Although there were twice as many repeat types in the gut than in saliva, the similar total number of spacers in the samples from each site suggested there were either longer CRISPR loci in organisms (on average) in saliva or there were more CRIPSR loci that shared the same repeat type among organisms in saliva.

We used the CRISPR spacers to detect matches in the NCBI database of viral genomes as well as across the data generated in this study. We found significantly more matches (and sequence similarity) to the data generated from our samples than to the NCBI virus database (Supplemental Fig. S12A). Within the metagenomic data, there were 78,054 total matches between spacers and scaffolds (Fig. 6B). On average, ∼32% of the spacers had at least one match to a scaffold of one of the body sites (vaginal: 31%; saliva: 38%; gut: 28%). Gut and saliva spacers had the most matches to scaffolds from the same body site. However, vaginal spacers had more matches to the gut scaffolds, both in terms of total matches between spacers and scaffolds (Fig. 6B) as well as in terms of the number of spacers with at least one match (Supplemental Fig. S12B). This was also true when vaginal spacers were viewed between and within individuals (Supplemental Table S3). It is possible that sequencing depth may have resulted in under-sampling of vaginal scaffold matches or that there were more scaffolds from the gut that serve as potential targets for vaginal spacers due to the higher diversity in the gut.

### Typical vaginal taxa are detected in the gut

While common gut *Lactobacillus* species such as *L. acidophilus, L. rhamnosus*, and *L. lactis* (Human Microbiome Project 2012) were identified in the metagenomic sequences assembled from gut communities, species typically seen in vaginal communities were found in 7 of the 10 subjects studied here. Near-complete genomes and shorter genomic fragments (3.5 -180 kb binned) of *L. iners* (in Term2, Term4, and Pre4), *L. crispatus* (in Term5, Term6 and Pre1), and *G. vaginalis* (in Pre2 and Pre4) were recovered from gut samples.

In all cases, the typical vaginal taxon found in the gut (at very low abundance) was the dominant bacterial species in the corresponding vaginal community of the same subject. Furthermore, genomic fragments from both *L. iners* and *G. vaginalis*, which were assembled from only one vaginal sample in subject Pre4 (Fig. 1) were detected in that subject’s gut. Sharing of *L. iners, L. crispatus*, and *G. vaginalis* between the vagina and gut was supported by an assessment of the presence of these vaginal bacteria in gut samples from the larger Stanford cohort. Of 41 subjects who provided both vaginal and gut samples (both stool and rectal swabs) (DiGiulio et al. 2015), 34 had 16S rRNA gene sequences of typical vaginal bacteria in their gut samples, and 33 had sequences from as many as 5 of their own vaginal species present in at least one of their gut samples (Supplemental Table S4). Evaluation of whole-genome core SNPs and clusters of orthologous genes in *L. iners* and *L. crispatus* reconstructed genome sequences indicated that the vaginal and gut sequences recovered within the same subject were more closely related to each other than to genome sequences assembled or identified in other subjects, suggesting that strains of vaginal *Lactobacillus* species from within the same subject come from the same ancestor (Supplemental Fig. S11A; Supplemental Fig. S12A).

The vaginal and gut genomes recovered here likely came from different populations, as evidenced by the amount of variation observed in full-genome and genomic fragment alignments within subjects, and the average pairwise nucleotide identity between the co-assembled genomes (Supplemental Fig. S13). Furthermore, SNPs within the vaginal read data mapping to the co-assembled genomes of *L. iners* and *L. crispatus* from the vagina indicated strain variation (*L. iners* and *L. crispatus* were deeply sequenced to a median of 219x coverage, with a range between 17x and 1180x). However, the frequency of SNPs unique to the gut read data that mapped to the corresponding vaginal strain genome from the same subject indicates that the pool of sequences used to assemble each of the genomes in the vagina and gut were not the same (Table 2, number of SNPs deemed significant based on the probability that a SNP is a product of random variation or sequencing error). For example, 1% of the SNPs that mapped to the vaginal *L. iners* genome from subject Term4 were unique to that subject’s gut read data (>400 SNPs distributed across the genome). Given that *L. iners* was assembled from the vagina of subject Term4 at 168x coverage, we would expect to recover almost all SNPs present in the gut read data (which achieved 7x coverage) if the sequences came from the same source.

**Table 2.**
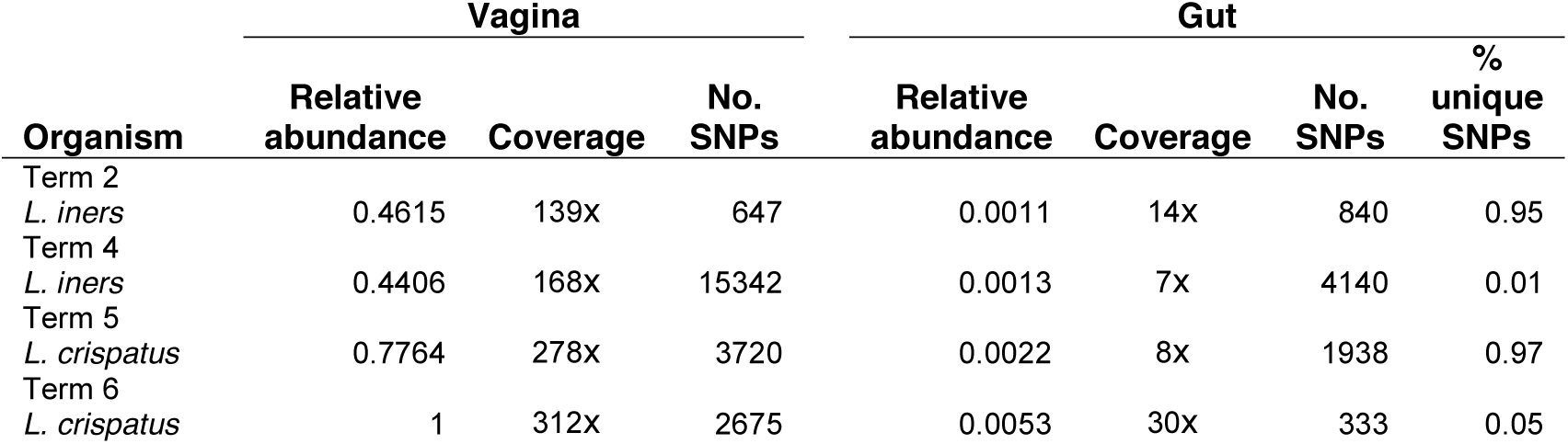
SNP analysis of the read data from vaginal and gut samples that mapped to the near-complete genomes of *L. iners* and *L. crispatus* recovered from the vagina. Sequencing reads from the vagina and gut of subjects Term2, Term4, Term5, and Term6 were mapped to the assembled genome of the dominant vaginal species for that same subject, and the occurrence of significant SNPs in the read data was determined using VarScan. The average relative abundance of each genome in the vagina and gut was estimated from the relative abundance of 16 single-copy ribosomal protein sequences, averaged across all samples sequenced from an individual subject. The percent SNPs unique to the gut genome was determined by comparing each SNP within a subject and filtering out significant SNPs that occurred at both body sites.

## DISCUSSION

The human vagina, oral cavity, and gut in combination with the microbial organisms, their genomes and functions within these body sites (microbiomes) reveal mutualistic relationships in which the host provides nutrients and a specialized niche, and the microbiota provides protection against pathogens and beneficial signals to the immune system, among other benefits (Fujimura et al. 2010; Ma et al. 2012). Here, we used whole community shotgun sequencing of human microbiome samples from three body sites (vagina, oral cavity, and gut) to characterize metagenome composition patterns over the course of pregnancy. High inter-individual variability and dynamic community composition across gestational age were observed at all body sites.

In the vagina, a direct relationship between gestational age and increasing taxonomic richness was observed in *L. iners*-dominated communities. This finding was also evident from re-evaluating the previously-published 16S rRNA gene amplicon data from the Stanford population (DiGiulio et al. 2015; Callahan et al. 2017). Although vaginal communities from subjects with abundant *Shuttleworthia* in the UAB cohort also showed statistically significant increases in taxonomic richness with gestational age, this association might be spurious due to the low sample size in *Shuttleworthia*-dominated communities. A significant gestation-associated increase in Shannon’s diversity index in *L. iners-* and *L. crispatus*-dominated communities from the Stanford, but not the UAB population, may be due to their different racial backgrounds, with a higher frequency of Caucasian and Asian individuals at Stanford, and a higher frequency of African American women at UAB (Callahan et al. 2017). Another difference is that, unlike at Stanford, women at UAB were treated with progesterone because of their history of prior preterm birth (Callahan et al. 2017). Although others have shown no effect of progesterone administration on microbiota composition over time (Kindinger et al. 2017), we cannot rule out an effect here.

L. crispatus has been correlated with stability over time in pregnancy, while *L. iners* has been associated with higher CST transition frequencies and instability (DiGiulio et al. 2015; Kindinger et al. 2017). *L. iners* is thought to be less competitive with *Gardnerella vaginalis*, and thus, to permit more frequent transitions to a high diversity community type (Ravel et al. 2011; Brooks et al. 2016; Callahan et al. 2017). Our data lend further support to the distinct ecology and less protective properties of this particular *Lactobacillus* species.

Analyses of community gene content indicated that while the individual was the strongest source of sample-to-sample variation, a significant gestational age (time) effect was observed for most subjects at all body sites. In addition, the dominant taxon (in vaginal communities) and adverse pregnancy outcome, e.g., preeclampsia (in gut and saliva communities) also appeared to be significant factors. For example, samples from subjects with a BMI > 25 appear to cluster closer to each other than to samples from other subjects. The small number of subjects in this study and the even smaller number with adverse pregnancy outcomes render any effort at this point to infer biological significance to this association problematic.

A previous analysis of the gut microbiota at two time points during pregnancy from a larger cohort of pregnant women reported an increase in diversity from the first to the third trimester (Koren et al. 2012). In contrast, our data indicated a decrease in diversity over time in our 10 subjects. Detailed longitudinal analyses of larger numbers of subjects, well-characterized for the presence or absence of gestational diabetes, BMI, and adverse outcomes will be needed for identification of biological leads.

The genomes of six strains of *Gardnerella vaginalis* were reconstructed and analyzed. Although *G. vaginalis* has been associated with bacterial vaginosis and with increased risk of preterm birth (Schwebke et al. 2014; DiGiulio et al. 2015; Callahan et al. 2017), it is also commonly found in vaginal CST IV communities of healthy women (Romero et al. 2014). *G. vaginalis* “genotypes” have been proposed previously: four main types were identified based on ∼473 concatenated core-gene alignments (Ahmed et al. 2012); three groups were suggested based on conserved chaperonin *cpn60* genes (Schellenberg et al. 2016); four genotypes were proposed with molecular genotyping (Santiago et al. 2011); and four clades were suggested from phylogenies reconstructed from core-gene sets (Malki et al. 2016). Phylogenomic trees reconstructed from full-genome alignments, as well as from core SNPs, based on 34 publicly available genomes along with the genomes recovered here suggested a fifth divergent clade within the *G. vaginalis* tree. We identified groups of genes enriched in specific functions within each clade. None of the publicly available genome sequences were generated from isolates from pregnant subjects: some came from asymptomatic healthy women, while other “pathogenic” strains were isolated from subjects with bacterial vaginosis (BV) (Yeoman et al. 2010). However, healthy vs. “pathogenic” strains did not map to specific clades. Consistent with previous comparative genomics reports of *G. vaginalis* (Ahmed et al. 2012; Malki et al. 2016), the *G. vaginalis* genomes recovered here show high degrees of re-arrangements with many unique islands. Although multiple *G. vaginalis* strains were isolated from the same subject previously (Ahmed et al. 2012), the co-occurring genomes were not compared, and the importance of co-occurring strains was not addressed. The results presented here suggest specialized functions encoded by closely related *G. vaginalis* strains and the possibility that different strains play different roles in health and disease.

CRISPR-Cas systems are intrinsically related to the mobile elements (such as phage and plasmid populations) that have targeted the genomic host in which they reside. Thus, CRISPR spacers allow an indirect assessment of the mobile element pool in the sampled habitat. The relative number of spacer matches in gut and saliva samples from this study indicate that phage populations are better adapted to their local environments than to environments found in other individuals. Previous findings have shown that samples from the same individual provide the highest number of matches for spacers from that same individual (Pride et al. 2011; Rho et al. 2012; Robles-Sikisaka et al. 2013; Abeles et al. 2014). However, we detected phage populations that are common enough between body sites and subjects to be targeted by the same CRISPR spacers, suggesting that similar mobile elements have been encountered by the microbial communities at different body sites and by different individuals.

Although many *Lactobacillus* species have been identified in the human gut, they represent relatively small proportions in 16S rRNA gene amplicon surveys (reviewed by Walters. 2017). *Lactobacillus crispatus, Lactobacillus iners, Lactobacillus gasseri,* and *Gardnerella vaginalis* are generally found in the vaginal (Human Microbiome Project. 2012; Romero et al. 2014; and reviewed by Hickey et al. 2012), although *L. crispatus* and *L. gasseri* have also been detected in the human gut (reviewed by Walter 2017), and other species, such as *L. rhamnosus* have been isolated from the vagina (Pascual 2008). Near-complete genomes and genomic fragments of *L. iners, L. crispatus,* and *G. vaginalis* were recovered from gut samples in this study, and the *Lactobacillus* species found in the gut of a specific subject was the dominant species found in the vagina of the same subject. Genome reconstruction of identical strains from samples of different body sites in the same subject has been reported previously based on metagenomic data from infants (Olm et al. 2017). One explanation for the finding of the same species in vaginal and gut samples of the same subject is contamination during the self-sampling procedure (likely when rectal swabs were provided in lieu of stool samples) or contamination during library preparation. However, given that the concurrent vaginal species occurred at high frequencies in both stool and rectal swab samples, and given the variation observed in the reconstructed genomes (from re-arrangements and genomic islands) and in the SNP patterns, we believe that the strains are not identical and are, therefore, likely not due to contamination. Colonization of the vagina and gut body sites is unlikely to occur independently (the same vaginal species was always found in the woman’s gut, with only one exception in the Stanford cohort). Colonization may occur from the mother at birth (especially if the subjects were delivered vaginally) or colonization from self at an earlier time point. Although there are no previous reports, to our knowledge, about shared bacterial species in the vagina and gut of the same individual, the phenomenon is unlikely to be limited to pregnant women. Examination of taxa reported in the HMP (Human Microbiome Project 2012) indicated the presence of *L. iners* and *Gardnerella* in gut samples from a few female subjects but in none of the male subjects. The paucity of evidence of *L. iners, L. crispatus*, and *G. vaginalis* in gut samples in the HMP study may be due to the difficulty in classifying short amplicon sequences at the species level, in addition to the shallow sequencing achieved in the HMP (Human Microbiome Project 2012).

The results reported here suggest dynamic behavior in the microbiome during pregnancy and highlight the importance of genome-resolved strain analyses to further our understanding of the establishment and evolution of the human microbiome.

## METHODS

### Sample Selection, DNA sample preparation and whole community metagenomic sequencing

A subset of subjects from the cohort examined with 16S rRNA gene amplicon analysis in (DiGiulio et al. 2015) were selected for this study. Six of these subjects (Term1-6) gave birth at term (after 37 weeks) while four of these subjects (Pre1-4) gave birth prematurely. Two of the term pregnancies and one preterm pregnancy were complicated by preeclampsia, one by oligohydramnios, and one by type 2 diabetes. Subjects and samples were selected based on whether samples were available at regular intervals (every three gestational weeks) from all 3 body sites, and on similar gestational dates across subjects. A total of 292 samples (101 vaginal swabs, 101 saliva, and 90 stool or rectal swabs) were selected for shotgun metagenomic analysis. No negative controls were performed for this study.

### Sequence processing and assembly

More than 7 billion paired-end reads were obtained, low quality bases were trimmed using Sickle with default parameters (Joshi and Fass 2011), adapter sequences were removed with SeqPrep (John JS 2011), and sequences >100 bp were retained. Trimmed reads were mapped to the human genome version GRCh37 with BWA (Li and Durbin 2009), and human reads were filtered out. Alignments were originally performed before release of Genome Reference Consortium Human Build 38 in December 2013. Regardless of the reference, some human derived reads usually remain after filtering due to natural variation in the human genome. We do not expect that there would be any differences in our conclusions if we were to have used Build 38, as the mapping across our datasets was done relative to the same human genome Build. Non-human reads were co-assembled across all time points per subject, per body site, with IDBA_UD (Peng et al. 2012) using default parameters. Scaffolds from vaginal samples were binned based on %GC and coverage using the public knowledgebase, ggKbase (http://ggkbase.berkeley.edu), as well as time-series information using ESOM (Dick et al. 2009). Genes were predicted with Prodigal (Hyatt et al. 2010), tRNAs with tRNA-scan (Lowe and Eddy 1997), and functional annotations were assigned from BLAST (Camacho et al. 2008) searches against the KEGG (Kanehisa and Goto 2000) and UniRef100 (The UniProt Consortium 2017) databases.

### Assessing the community composition

Full-length 16S rRNA gene sequences were reconstructed per body site with EMIRGE (Miller et al. 2011), using the SILVA database SSURef 111 (Pruesse et al. 2007) and run with 100 iterations. Sequences that were identified as chimeras with any two of the three chimera check software packages, UCHIME (Edgar et al. 2011), DECIPHER (Wright et al. 2012), and Bellerophon (Huber et al. 2004), were removed. The full-length 16S rRNA gene sequences from the six *G. vaginalis* strain genomes, and those from 22 publicly available genomes were aligned with MUSCLE (Edgar 2004) and visualized with Geneious (Kearse et al. 2012). 16S rRNA gene amplicon data from our previous work (DiGiulio et al. 2015; Callahan et al. 2017) were used to evaluate richness trends with gestation.

### Calculations of relative abundances of genes and genome bins

Non-human reads were mapped to the co-assembled scaffolds from each subject (per sample) using Bowtie 2 (Langmead and Salzberg 2012) with default parameters. Vaginal genome bin abundance was calculated by dividing the numbers of reads that mapped to all scaffolds within a bin, by the total number of reads that mapped to all scaffolds within a subject.

16 single-copy ribosomal protein (RP) gene sets were used to estimate bacterial diversity within body sites, as previously described (Hug et al. 2016). In total, 22,737 ribosomal proteins annotated as L14b/L23e, L15, L16/L10E, L18P/L5E, L2 rplB, L22, L24, L3, L4/L1e, L5, L6, S10, S17, S19, S3, and S8 were identified in 4,650 scaffolds from all body sites. Individual RP gene sets were clustered at 99% average amino acid identity (AAI) with usearch64 (Edgar 2010), and scaffolds containing the same sets of clustered RPs were considered to be derived from the same taxon (accounting for a total of 4,024 taxa). Scaffold coverage was used to estimate read counts for predicted genes, UniRef90 gene families, and RP abundance tables (taxa abundance tables)

### Statistical analysis

NMDS ordination was performed on Bray-Curtis distance matrices of variance-stabilizing transformed UniRef90 gene family abundance tables using the DESeq2 (Love et al. 2014) and phyloseq (McMurdie and Holmes 2013) packages in R (R Core Team 2017). RDA was performed directly on variance-stabilizing transformed UniRef90 gene family abundance tables using the same R packages used for NMDS, with the formula: genes ∼ gestational age + complication (for gut and oral samples), and genes ∼ gestational age + top taxon (vaginal samples). The 16S rRNA gene tables, UniRef90 gene family tables, RP tables, pathway abundance tables, and R code are available at the Stanford Digital Repository at https://purl.stanford.edu/vp282bh3698 (.rmd files contain the code,.RData files contain data tables).

### Functional pathway presence and abundance analysis

Functional pathways were predicted with HUMAnN2 (Abubucker et al. 2012) using the read FASTQ files. Pathway abundance tables were variance-stabilized with DESeq2 in R and are available in the.RData files at the Stanford Digital Repository at https://purl.stanford.edu/vp282bh3698.

### G. vaginalis, L. iners, and *L. crispatus phylogenetic trees and genome comparisons*

34 publicly available *G. vaginalis* strain genomes, plus the six *G. vaginalis* genomes recovered in this study were aligned with progressiveMauve (Darling et al. 2010). Similarly, six *L. iners* genomes assembled from vaginal samples, the two from gut samples, and 16 publicly available *L. iners* genomes were aligned with progressiveMauve, with *L. iners* DSM 13335 as a reference. In addition, 37 publicly available *L. crispatus* genomes (assembled in < 120 scaffolds), as well as six *L. crispatus* genomes reconstructed from vaginal and gut samples, were aligned with progressiveMauve, with the *L. crispatus ST1* representative genome as a reference. The output of progressiveMauve was visualized and exported in standard xmfa format using Gingr (Treangen et al. 2014), and converted to FASTA format using a publicly available Perl script (Jolley K 2010). For *G. vaginalis*, the multiple genome alignment in FASTA format was used to construct the phylogenetic trees using FastTree (Price et al. 2009) version 2.1.8 with parameters –nt –gtr –gamma, and visualized using Geneious (Kearse et al. 2012). Orthologous gene groups between *G. vaginalis, L. iners* and *L. crispatus* genomes were exported from the Mauve alignments based on at least 60% identity over 70% alignment coverage. Orthologous groups were classified into functional categories represented in the Gene Ontology (GO) Database by mapping GI gene numbers to GO IDs in bioDBnet (Mudunuri et al. 2009). Hierarchical clustering on the Jaccard distances of the presence/absence orthologous groups was performed with the vegan package (Jari Oksanen et al. 2017), and the heatmap was created with the pheatmap package (Kolde 2015) in R (R Core Team 2017). The number of clusters was determined visually from the hierarchical clustering, and the list of genes within each cluster was obtained with the function cutree from the R package “stats” (R Core Team. 2017). The enrichment of genes within functional categories was estimated from the number of genes in each functional category within each cluster divided by the total number of orthologous genes in each cluster.

kSNP3.0 (Gardner et al. 2015) was used to build phylogenetic trees from whole-genome SNP patterns in *Lactobacillus iners, Lactobacillus crispatus*, and *Gardnerella vaginalis*. iRep (with default parameters) was used to estimate the replication rate of genomes reconstructed from the vaginal metagenomic datasets as in Brown *et al*. (Brown et al. 2015).

### Strain comparison across body sites

Bowtie 2 (Langmead and Salzberg 2012) was used to identify reads from the gut dataset that mapped to the genome of each *Lactobacillus* species assembled from the gut. Then Bowtie 2 was used to map that subset of reads against the corresponding vaginal population genome assembled from the same subject. Reads from the vaginal samples were also mapped to the corresponding *Lactobacillus* species assembled from the vagina. VarScan (Koboldt et al. 2012) was used to identify the SNPs, with parameters, minimum coverage > 3 and p value < 0.05.

### Identification of CRISPR repeats/spacers, predicted Cas proteins, and CRISPR-Cas systems

The program Crass (Skennerton et al. 2013), with default settings, was used to identify CRISPR repeats and spacers from all reads from each sample. The output was parsed using CRISPRtools (http://ctskennerton.github.io/crisprtools/). HMMER3 (Mistry et al. 2013) was used to search all predicted proteins against a Cas proteins HMM database created from alignments from refs (Makarova et al. 2015; Shmakov et al. 2015). The HMM database can be found in Supp. Data 6 from (Burstein et al. 2016). CRISPR-Cas system types were identified based on the presence of 2 predicted Cas proteins. CRISPR-Cas types were only calculated for gut and saliva samples (vaginal samples had low organismal diversity). Cas types that were incomplete were marked as unknown/unclassified.

### Detection of potential phage and plasmid sequences and dynamics of spacer matching

For body site-specific analysis, CRISPR repeats and spacers were extracted from the reads combined across all sites. For individual analyses, CRISPR repeats and spacers were combined for each body site within subjects (term1 vaginal reads, term 1 gut reads, term1 saliva reads, etc.). A spacer was recorded as a match to a scaffold if the spacer matched with at least 85% identity across 85% of the spacer length against a scaffold using BLASTn (Camacho et al. 2008), with the parameter “-task blastn-short”.

## DATA ACCESS

The generated sequencing reads have been deposited to the NCBI Sequencing Read Archive (SRA, https://www.ncbi.nlm.nih.gov/sra) under BioProject accession number PRJNA288562. The specific BioSample accession numbers (292 in total) are listed in Supplemental Table S5. Assemblies, protein sequences, gbk files, gene and protein abundance tables, metadata, and R code are publicly available at the Stanford Digital Repository at https://purl.stanford.edu/vp282bh3698. R markdown and data files are available as Supplemental Data. The 29 genome assemblies are available at GenBank Genomes (accession numbers pending).

## ACKNOWLEDGMENTS

We are grateful to study participants, as well as Cele Quaintance, Nick Scalfone, Ana Laborde, March of Dimes Prematurity Research Center study coordinators, and nursing staff in the obstetrical clinics and the labor and delivery unit of Lucille Packard Children’s Hospital. We thank Alvaro Hernandez at the University of Illinois Roy J. Carver Biotechnology Center. We thank David Burstein for advice and scripts for Cas protein analysis, and Elizabeth Costello for helpful discussion. This research was supported by the March of Dimes Prematurity Research Center at Stanford University School of Medicine, the Thomas C. and Joan M. Merigan Endowment at Stanford University (D.A.R.), the Chan Zuckerburg Biohub (D.A.R.), and the Eunice Kennedy Shriver National Institute Of Child Health & Human Development of the National Institutes of Health under Award K99HD090290 (D.S.A.G.).

## REFERENCES

Abeles SR, Robles-Sikisaka R, Ly M, Lum AG, Salzman J, Boehm TK, Pride DT. 2014. Human oral viruses are personal, persistent and gender-consistent. The ISME journal 8: 1753–15 1767.

Abubucker S, Segata N, Goll J, Schubert AM, Izard J, Cantarel BL, Rodriguez-Mueller B, Zucker J, Thiagarajan M, Henrissat B et al. 2012. Metabolic reconstruction for metagenomic data and its application to the human microbiome. PLoS Comput Biol 8: e1002358.

Ahmed A, Earl J, Retchless A, Hillier SL, Rabe LK, Cherpes TL, Powell E, Janto B, Eutsey R, Hiller NL et al. 2012. Comparative genomic analyses of 17 clinical isolates of Gardnerella vaginalis provide evidence of multiple genetically isolated clades consistent with subspeciation into genovars. J Bacteriol 194: 3922–3937.

Andersson AF, Banfield JF. 2008. Virus population dynamics and acquired virus resistance in 25 natural microbial communities. Science 320: 1047–1050.

Bolotin A, Quinquis B, Sorokin A, Ehrlich SD. 2005. Clustered regularly interspaced short palindrome repeats (CRISPRs) have spacers of extrachromosomal origin. Microbiology 151: 2551–2561.

Brooks JP, Buck GA, Chen G, Diao L, Edwards DJ, Fettweis JM, Huzurbazar S, Rakitin A, Satten GA, Smirnova E, Waks Z, Wright ML, Yanover C, Zhou YH. 2017. Changes in vaginal community state types reflect major shifts in the microbiome. Microb Ecol Health Dis 28: 1303265.

Brown CT, Hug LA, Thomas BC, Sharon I, Castelle CJ, Singh A, Wilkins MJ, Wrighton KC, Williams KH, Banfield JF. 2015. Unusual biology across a group comprising more than 10 15% of domain Bacteria. Nature 523: 208–211.

Burstein D, Sun CL, Brown CT, Sharon I, Anantharaman K, Probst AJ, Thomas BC, Banfield JF. 2016. Major bacterial lineages are essentially devoid of CRISPR-Cas viral defence systems. Nature communications 7: 10613.

Callahan BJ, DiGiulio DB, Goltsman DSA, Sun CL, Costello EK, Jeganathan P, Biggio JR, Wong RJ, Druzin ML, Shaw GM et al. 2017. Replication and refinement of a vaginal microbial signature of preterm birth in two racially distinct cohorts of US women. Proceedings of the National Academy of Sciences of the United States of America 114: 9966–9971.

Camacho C, Coulouris G, Avagyan V, Ma N, Papadopoulos J, Bealer K, Madden TL. 2008 BLAST+: architecture and applications. BMC Bioinformatics 10:421

Charbonneau MR, Blanton LV, DiGiulio DB, Relman DA, Lebrilla CB, Mills DA, Gordon JI. 2016. A microbial perspective of human developmental biology. Nature 535: 48–55.

Cho I, Blaser MJ. 2012. The human microbiome: at the interface of health and disease. Nature reviews Genetics 13: 260–270.

Costello EK, Stagaman K, Dethlefsen L, Bohannan BJ, Relman DA. 2012. The application of ecological theory toward an understanding of the human microbiome. Science 336: 1255–1262.

Darling AE, Mau B, Perna NT. 2010. progressiveMauve: multiple genome alignment with gene gain, loss and rearrangement. PloS one 5: e11147.

Dick GJ, Andersson AF, Baker BJ, Simmons SL, Thomas BC, Yelton AP, Banfield JF. 2009. Community-wide analysis of microbial genome sequence signatures. Genome Biol 10: R85.

DiGiulio DB, Callahan BJ, McMurdie PJ, Costello EK, Lyell DJ, Robaczewska A, Sun CL, Goltsman DSA, Wong RJ, Shaw G et al. 2015. Temporal and Spatial Variation of the Human Microbiota During Pregnancy. Proc Natl Acad Sci USA 112:11060–5.

Edgar RC. 2004. MUSCLE: multiple sequence alignment with high accuracy and high throughput. Nucleic Acids Res 32: 1792–1797.

Edgar RC. 2010. Search and clustering orders of magnitude faster than BLAST. Bioinformatics 26: 2460–2461.

Edgar RC, Haas BJ, Clemente JC, Quince C, Knight R. 2011. UCHIME improves sensitivity and speed of chimera detection. Bioinformatics 27: 2194–2200.

Fujimura KE, Slusher NA, Cabana MD, Lynch SV. 2010. Role of the gut microbiota in defining human health. Expert review of anti-infective therapy 8: 435–454.

Gajer P, Brotman RM, Bai G, Sakamoto J, Schutte UM, Zhong X, Koenig SS, Fu L, Ma ZS, Zhou X et al. 2012. Temporal dynamics of the human vaginal microbiota. Science translational medicine 4: 132ra152.

Gardner SN, Slezak T, Hall BG. 2015. kSNP3.0: SNP detection and phylogenetic analysis of genomes without genome alignment or reference genome. Bioinformatics 31: 2877–2878.

Garrett WS, Gordon JI, Glimcher LH. 2010. Homeostasis and inflammation in the intestine. Cell 140: 859–870.

Hickey RJ, Zhou X, Pierson JD, Ravel J, Forney LJ. 2012. Understanding vaginal microbiome complexity from an ecological perspective. Translational research: the journal of laboratory and clinical medicine 160: 267–282.

Hillier SL, Nugent RP, Eschenbach DA, Krohn MA, Gibbs RS, Martin DH, Cotch MF, Edelman R, Pastorek JG, 2nd, Rao AV et al. 1995. Association between bacterial vaginosis and preterm delivery of a low-birth-weight infant. The Vaginal Infections and Prematurity Study Group. The New England journal of medicine 333: 1737–1742.

Horvath P, Barrangou R. 2010. CRISPR/Cas, the immune system of bacteria and archaea. Science 327: 167–170.

Huber T, Faulkner G, Hugenholtz P. 2004. Bellerophon: a program to detect chimeric sequences in multiple sequence alignments. Bioinformatics 20: 2317–2319.

Hug LA, Baker BJ, Anantharaman K, Brown CT, Probst AJ, Castelle CJ, Butterfield CN, Hernsdorf AW, Amano Y, Ise K et al. 2016. A new view of the tree of life. Nat Microbiol 1: 16048.

Human Microbiome Project C. 2012. Structure, function and diversity of the healthy human microbiome. Nature 486: 207–214.

Hyatt D, Chen GL, Locascio PF, Land ML, Larimer FW, Hauser LJ. 2010. Prodigal: prokaryotic gene recognition and translation initiation site identification. BMC bioinformatics 11: 119.

Hyman RW, Fukushima M, Jiang H, Fung E, Rand L, Johnson B, Vo KC, Caughey AB, Hilton JF, Davis RW et al. 2014. Diversity of the vaginal microbiome correlates with preterm birth. Reproductive sciences 21: 32–40.

Jari Oksanen FGB, Michael Friendly, Roeland Kindt, Pierre Legendre DM, Peter R. Minchin, R. B. O’Hara, Gavin L., Simpson PS, M. Henry H. Stevens, Eduard Szoecs and Helene, Wagner. 2017. vegan: Community Ecology Package.

John JS. 2011. SeqPrep. https://github.com/jstjohn/SeqPrep.

Jolley K. 2010. xmfa2fasta.pl. https://github.com/kjolley/seq_scripts/blob/master/xmfa2fasta.pl

Joshi NA, Fass JN. 2011. Sickle: A sliding-window, adaptive, quality-based trimming tool for FastQ files (Version 1.33). https://github.com/najoshi/sickle.

Kanehisa M, Goto S. 2000. KEGG: kyoto encyclopedia of genes and genomes. Nucleic Acids Res 28: 27–30.

Karginov FV, Hannon GJ. 2010. The CRISPR system: small RNA-guided defense in bacteria and archaea. Mol Cell 37: 7–19.

Kearse M, Moir R, Wilson A, Stones-Havas S, Cheung M, Sturrock S, Buxton S, Cooper A, Markowitz S, Duran C et al. 2012. Geneious Basic: an integrated and extendable desktop software platform for the organization and analysis of sequence data. Bioinformatics 28: 1647–1649.

Kindinger LM, Bennett PR, Lee YS, Marchesi JR, Smith A, Cacciatore S, Holmes E, Nicholson JK, Teoh TG, MacIntyre DA. 2017. The interaction between vaginal microbiota, cervical length, and vaginal progesterone treatment for preterm birth risk. Microbiome 19;5(1):6.

Koboldt DC, Zhang Q, Larson DE, Shen D, McLellan MD, Lin L, Miller CA, Mardis ER, Ding L, Wilson RK. 2012. VarScan 2: somatic mutation and copy number alteration discovery in cancer by exome sequencing. Genome Res 22:568–76.

Kolde R. 2015. pheatmap: Pretty Heatmaps.

Koren O, Goodrich JK, Cullender TC, Spor A, Laitinen K, Backhed HK, Gonzalez A, Werner JJ, Angenent LT, Knight R et al. 2012. Host remodeling of the gut microbiome and metabolic changes during pregnancy. Cell 150: 470–480.

Langmead B, Salzberg S. 2012. Fast gapped-read alignment with Bowtie 2. Nature Methods 9:357–359.

Li H, Handsaker B, Wysoker A, Fennell T, Ruan J, Homer N, Marth G, Abecasis G, Durbin R, 20 and 1000 Genome Project Data Processing Subgroup. 2009. The Sequence alignment/map (SAM) format and SAMtools. Bioinformatics 25: 2078–9

Li H, Durbin R. 2009. Fast and accurate short read alignment with Burrows-Wheeler transform. Bioinformatics 25: 1754–1760.

Love MI, Huber W, Anders S. 2014. Moderated estimation of fold change and dispersion for RNA-seq data with DESeq2. Genome Biol 15: 550.

Lowe TM, Eddy SR. 1997. tRNAscan-SE: a program for improved detection of transfer RNA genes in genomic sequence. Nucleic Acids Res 25: 955–964.

Ma B, Forney LJ, Ravel J. 2012. Vaginal microbiome: rethinking health and disease. Annual review of microbiology 66: 371–389.

Macklaim JM, Gloor GB, Anukam KC, Cribby S, Reid G. 2011. At the crossroads of vaginal health and disease, the genome sequence of Lactobacillus iners AB-1. Proceedings of the National Academy of Sciences of the United States of America 108 Suppl 1: 4688–4695.

Makarova KS, Wolf YI, Alkhnbashi OS, Costa F, Shah SA, Saunders SJ, Barrangou R, Brouns SJ, Charpentier E, Haft DH et al. 2015. An updated evolutionary classification of CRISPR-Cas systems. Nature reviews Microbiology 13: 722–736.

Malki K, Shapiro JW, Price TK, Hilt EE, Thomas-White K, Sircar T, Rosenfeld AB, Kuffel G, Zilliox MJ, Wolfe AJ et al. 2016. Genomes of Gardnerella Strains Reveal an Abundance of Prophages within the Bladder Microbiome. PloS one 11: e0166757.

Marraffini LA, Sontheimer EJ. 2010. CRISPR interference: RNA-directed adaptive immunity in bacteria and archaea. Nature reviews Genetics 11: 181–190.

MacIntyre DA, Chandiramani M, Lee YS, Kindinger L, Smith A, Angelopoulos N, Lehne B, Arulkumaran S, Brown R, Teoh TG, Holmes E, Nicoholson JK, Marchesi JR, Bennett PR. 2015. The vaginal microbiome during pregnancy and the postpartum period in a European population. Sci Rep 11;5:8988.

McMurdie PJ, Holmes S. 2013. phyloseq: an R package for reproducible interactive analysis and graphics of microbiome census data. PloS one 8: e61217.

Miller CS, Baker BJ, Thomas BC, Singer SW, Banfield JF. 2011. EMIRGE: reconstruction of full-length ribosomal genes from microbial community short read sequencing data. Genome Biol 12: R44.

Mistry J, Finn RD, Eddy SR, Bateman A, Punta M. 2013. Challenges in Homology Search: HMMER3 and Convergent Evolution of Coiled-Coil Regions. Nucleic Acids Res 41:e121.

Mojica FJ, Diez-Villasenor C, Garcia-Martinez J, Soria E. 2005. Intervening sequences of regularly spaced prokaryotic repeats derive from foreign genetic elements. J Mol Evol 60: 174–182.

Moriya Y, Itoh M, Okuda S, Yoshizawa AC, Kanehisa M. 2007. KAAS: an automatic genome annotation and pathway reconstruction server. Nucleic Acids Res 35: W182–185.

Mudunuri U, Che A, Yi M, Stephens RM. 2009. bioDBnet: the biological database network. Bioinformatics 25: 555–556.

Nuriel-Ohayon M, Neuman H, Koren O. 2016. Microbial Changes during Pregnancy, Birth, and Infancy. Front Microbiol 7: 1031.

Ojala T, Kankainen M, Castro J, Cerca N, Edelman S, Westerlund-Wikstrom B, Paulin L, Holm L, Auvinen P. 2014. Comparative genomics of Lactobacillus crispatus suggests novel mechanisms for the competitive exclusion of Gardnerella vaginalis. BMC Genomics 15: 1070

Olm MR, Brown CT, Brooks B, Firek B, Baker R, Burstein D, Soenjoyo K, Thomas BC, Morowitz M, Banfield JF. 2017. Identical bacterial populations colonize premature infant gut, skin, and oral microbiomes and exhibit different in situ growth rates. Genome Research 27: 601–612.

Parks DH, Imelfort M, Skennerton CT, Hugenholtz P, Tyson GW. 2015. CheckM: assessing the quality of microbial genomes recovered from isolates, single cells, and metagenomes. Genome Research 25: 1043–1055.

Peng Y, Leung HC, Yiu SM, Chin FY. 2012. IDBA-UD: a de novo assembler for single-cell and metagenomic sequencing data with highly uneven depth. Bioinformatics 28: 1420–1428.

Pourcel C, Salvignol G, Vergnaud G. 2005. CRISPR elements in Yersinia pestis acquire new repeats by preferential uptake of bacteriophage DNA, and provide additional tools for evolutionary studies. Microbiology 151: 653–663.

Price MN, Dehal PS, Arkin AP. 2009. FastTree: computing large minimum evolution trees with profiles instead of a distance matrix. Mol Biol Evol 26: 1641–1650.

Pride DT, Sun CL, Salzman J, Rao N, Loomer P, Armitage GC, Banfield JF, Relman DA. 2011. Analysis of streptococcal CRISPRs from human saliva reveals substantial sequence diversity within and between subjects over time. Genome research 21: 126–136.

Pruesse E, Quast C, Knittel K, Fuchs BM, Ludwig W, Peplies J, Glockner FO. 2007. SILVA: a comprehensive online resource for quality checked and aligned ribosomal RNA sequence data compatible with ARB. Nucleic Acids Res 35: 7188–7196.

R Core Team. 2017. R: A language and environment for statistical computing. R Foundation for Statistical Computing, Vienna, Austria. https://www.R-project.org/.

Ravel J, Gajer P, Abdo Z, Schneider GM, Koenig SS, McCulle SL, Karlebach S, Gorle R, Russell J, Tacket CO et al. 2011. Vaginal microbiome of reproductive-age women. Proceedings of the National Academy of Sciences of the United States of America 108 Suppl 1: 4680–4687.

Raymond KN, Dertz EA, Kim SS. 2003. Enterobactin: an archetype for microbial iron transport. Proceedings of the National Academy of Sciences of the United States of America 100: 3584–3588.

Rho M, Wu YW, Tang H, Doak TG, Ye Y. 2012. Diverse CRISPRs evolving in human microbiomes. PLoS Genet 8: e1002441.

Robles-Sikisaka R, Ly M, Boehm T, Naidu M, Salzman J, Pride DT. 2013. Association between living environment and human oral viral ecology. ISME Journal 7: 1710–1724.

Romero R, Hassan SS, Gajer P, Tarca AL, Fadrosh DW, Nikita L, Galuppi M, Lamont RF, Chaemsaithong P, Miranda J et al. 2014. The composition and stability of the vaginal microbiota of normal pregnant women is different from that of non-pregnant women. Microbiome 2: 4.

Santiago GL, Deschaght P, El Aila N, Kiama TN, Verstraelen H, Jefferson KK, Temmerman M, Vaneechoutte M. 2011. Gardnerella vaginalis comprises three distinct genotypes of which only two produce sialidase. American journal of obstetrics and gynecology 204: 450 e451–457.

Schellenberg JJ, Paramel Jayaprakash T, Withana Gamage N, Patterson MH, Vaneechoutte M, Hill JE. 2016. Gardnerella vaginalis Subgroups Defined by cpn60 Sequencing and Sialidase Activity in Isolates from Canada, Belgium and Kenya. PloS one 11: e0146510.

Schwebke JR, Muzny CA, Josey WE. 2014. Role of Gardnerella vaginalis in the pathogenesis of bacterial vaginosis: a conceptual model. Journal of infectious diseases 210: 338–343.

Skennerton CT, Imelfort M, Tyson GW. 2013. Crass: identification and reconstruction of CRISPR from unassembled metagenomic data. Nucleic Acids Res 41: e105.

Smith SB, Ravel J. 2017. The vaginal microbiota, host defence and reproductive physiology. J Physiol 595: 451–463.

Stout MJ, Zhou Y, Wylie KM, Tarr PI, Macones GA, Tuuli MG. 2017. Early pregnancy vaginal microbiome trends and preterm birth. Am J Obstet Gynecol. 217(3):356.e1-356.e18.

Sun CL, Thomas BC, Barrangou R, Banfield JF. 2016. Metagenomic reconstructions of bacterial CRISPR loci constrain population histories. ISME Journal 10: 858–870.

Tabatabaei N, Eren AM, Barreiro LB, Yotova V, Dumaine A, Allard C, Fraser WD. 2018. Vaginal microbiome in early pregnancy and subsequent risk of spontaneous preterm birth: a case-control study. BJOG. 23. [Epub ahead of print]

The UniProt Consortium. 2017. UniProt: the universal protein knowledgebase. Nucleic Acids Research 45: D158–D169

Treangen TJ, Ondov BD, Koren S, Phillippy AM. 2014. The Harvest suite for rapid core-genome alignment and visualization of thousands of intraspecific microbial genomes. Genome Biol 15: 524.

Walter J. Ecological role of lactobacilli in the gastrointestinal tract: implications for fundamental and biomedical research. Appl Environ Microbiol 74(16):4985–96.

Wright ES, Yilmaz LS, Noguera DR. 2012. DECIPHER, a search-based approach to chimera identification for 16S rRNA sequences. Aplied and environmental microbiology 78: 717–725.

Yeoman CJ, Yildirim S, Thomas SM, Durkin AS, Torralba M, Sutton G, Buhay CJ, Ding Y, Dugan-Rocha SP, Muzny DM et al. 2010. Comparative genomics of Gardnerella vaginalis strains reveals substantial differences in metabolic and virulence potential. PloS one 5: e12411.

Zhou X, Brown CJ, Abdo Z, Davis CC, Hansmann MA, Joyce P, Foster JA, Forney LJ. 2007. Differences in the composition of vaginal microbial communities found in healthy Caucasian and black women. ISME Journal 1: 121–133.

